# ADP-MoA: a platform for screening antibiotic activity and their mechanism of action in *Pseudomonas aeruginosa*

**DOI:** 10.1101/2024.11.08.622684

**Authors:** Estela Ynés Valencia Morante, Viviane Abreu Nunes, Felipe S Chambergo, Beny Spira

**Author notes:** Address correspondence to Beny Spira,.

## Abstract

The emergence and proliferation of multidrug-resistant bacteria pose a major threat to global public health. To address an imminent crisis, it is essential to identify and characterize new antibacterial molecules. With that in mind, we developed the ADP-MoA platform, that facilitates the discovery of new antibiotics and provides preliminary insights into their mechanisms of action. The basic idea is to simultaneously visualize antibiotic activity – growth inhibition, along with one of the three classic antibiotics mechanisms of action: DNA damage/inhibition of DNA replication, protein synthesis inhibition and cell wall damage. The platform consists of three different chromosomal fusions between the promoters of *recA, ampC* or *armZ* and the *luxCDABE* operon. The platform was constructed and hitherto tested in the pathogenic opportunistic bacterium *Pseudomonas aeruginosa*. As a proof of concept we showed that the promoter fusions were each activated by the expected antibiotics with known mechanisms of action. The *armZ*::*luxCDABE* fusion responded to antibiotics that inhibit protein synthesis (macrolides, chloramphenicol, tetracyclines and aminoglycosides), *ampC*::*luxCDABE* was induced by β-lactams and *recA*::*luxCDABE* was induced by quinolones. Interestingly, ciprofloxacin induced P*ampC* and P*armZ* as well, albeit at a lower level. The ADP-MoA platform offers a readily implementable, low-cost approach with significant potential for high-throughput screening of antimicrobials against *P. aeruginosa* and other bacterial species.

## INTRODUCTION

Multi-resistant pathogenic bacteria are one of the main threats to public health worldwide. An estimated 1.27 million deaths occur annually due to pathogenic bacteria resistant to available antibiotics [1] and that in 2050 they will cause 10 million deaths per year, making them more lethal than cancer. Although antibiotic resistance concerns all pathogenic bacteria, the WHO Bacterial Priority Pathogens List [2] named 15 families of antibiotic-resistant pathogens, which have a strong negative impact on human and animal health. Among them is the opportunistic bacterium *Pseudomonas aeruginosa*. In particular, *P. aeruginosa* strain PA14 (UCBPP-PA14), isolated from a patient burn wound [3], presents an arsenal of molecular mechanisms that confer intrinsic and acquired resistance to multiple classes of antibiotics and many virulence genes, mostly related to the secretion of toxins and biofilm formation [4, 5].

The research and development of novel antibiotics by pharmaceutical companies has drastically decreased in recent years [6]. Preclinical antibiotic research (extraction, identification, functional analysis, molecular target validation, physicochemical characteristics, pharmacokinetic and pharmacodynamic studies) take approximately five years [7]. The simultaneous assessment of antibacterial activity and characterization of the mechanisms underlying this biological activity could shorten the pace of new antibiotics’ discovery [8].

In the present study, we report on the design and development of a bacterial genetic platform that provides a functional assay aimed at screening antimicrobial molecules (through growth inhibition) while simultaneously determining their mechanism of action (DNA damage/replication inhibition, ribosome inhibition, or cell wall damage). The set of biosensors that form the platform was named ADP-MoA, which stands for “Antibiotic discovery platform by mechanism of action”. At this stage, the platform was implemented in strain PA14 of *P. aeruginosa*, into which biosensors carrying specific promoters (*recA, ampC* or *armZ*) were fused to the *luxCDABE* operon (*lux* for the sake of brevity). These promoters were chosen based on their response to the presence of specific antibiotics with known mechanisms of action. The *recA* gene encodes a protein crucial for DNA repair and maintenance as part of the bacterial SOS response, with its transcription being induced by DNA damage or inhibition of DNA replication [9, 10]; *ampC* encodes a β-lactamase that is activated, via AmpR, in response to cell wall damage [11, 12] and *armZ* responds to ribosome-targeting antimicrobials [13, 14]. As a proof of concept, the three biosensors were tested with dozens of known antibiotics, validating their effectiveness and specificity.

## MATERIALS AND METHODS

### Bacterial strains and growth conditions

*P. aeruginosa* PA14 strain was used to host a chromosomal copy of each biosensor. *Escherichia coli* strains DH5α or DH10B were used for DNA manipulation. Bacteria were grown in lysogeny broth (LB) [15] at 37°C. When necessary, the medium was supplemented with 30 µg/mL gentamicin or 100 µg/mL ampicillin. Muller Hinton (MH) or MH-Agar media were used in the antimicrobial tests.

### Construction of the biosensors

The genomic DNA of strain PA14 was extracted using the Wizard™ Genomic DNA Purification Kit (Promega). DNA fragments corresponding to specific promoter regions were amplified by PCR with the help of the TransStart™ FastPfu DNA Polymerase kit (TransGen Biotech) and specific primers. The following amplicons were obtained (numbers correspond to the first and last base on the *P. aeruginosa* PA14 genome– GenBank number: CP034244.1): *recA* (1504655 to 1505155); *ampC* (933654 to 933962) and *armZ* (6432940 to 6432561). The amplicons were cloned in the BamHI/PstI sites of plasmid pUC18T-mini-Tn7T-*lux*-Gm [16] located upstream to the promoterless *lux* operon.

Following ligation, the plasmids were electrotransformed in *E. coli* strains DH10B or DH5α, and then electrotransformed in strain PA14 together with the helper plasmid pTNS3 that carries the genes encoding the TnsABCD site-specific transposition proteins [17]. Integration of the promoter-*lux* fusions 25 nucleotides downstream of the *glmS* gene in PA14 chromosome was verified by PCR, with primers PTn7R (5’-CACAGCATAACTGGACTGATTTC-3’) and PglmS-down (5’-GCACATCGGCGACGTGCTCTC-3’), as recommended by [17].

### Antibacterial activity and luminescence detection on plates

The Kirby-Bauer disc diffusion method [18] was used to test both antibacterial (growth inhibition) and biosensor activity in the presence of known antibiotics. Thirty eight different commercial disks (CECON, São Paulo-Brazil) were tested: amikacin (AMI, 30 µg), ampicillin (AMP, 10 µg), ampicillin + sulbactam (SBA, 20 µg), azithromycin (AZI, 15 µg), bacitracin (BAC, 10 µg), cefepime (CPM, 30 µg), cefoxitin (CFO, 30 µg), ceftriaxone (CRO, 30 µg), cefuroxime (CRX, 30 µg), chloramphenicol (CLO, 30 µg), Ciprofloxacin (CIP, 5 µg), clavulanic acid+ amoxicillin (AMC, 30 µg), clindamycin (CLI, 2 µg), doxycycline (DOX, 30 µg), enrofloxacin (ENO, 5µg), ertapenem (ETP, 10 µg), erythromycin (ERI, 15 µg), fosfomycin (FOS, 200 µg), gentamicin (GM, 10 µg), imipinem (IPM, 10 µg), kanamycin (CAN, 30 µg), levofloxacin (LVX, 5 µg), meropenem (MPM, 10 µg), nalidixic acid (NAL, 30), neomycin (NEO, 30 µg), nitrofurantoin (NIT, 300 µg), norfloxacin, (NOR, 10 µg), ofloxacin (OFX, 5 µg), penicillin (PEN, 10 U), piperacillin + tazobactam (PPT, 110 µg), rifampicin (RIF, 5 µg), streptomycin (EST, 10 µg), sulfadiazine + trimethoprim (SDT, 25 µg), sulfamethoxazole + trimethoprim (SUT, 25 µg), tetracycline (TET, 30 µg), ticarcillin (TIC, 75 µg), tobramycin (TOB, 10 µg) and trimethoprim (TRI, 5 µg). The disks were placed on MH agar plates, previously spread with 0.5 McFarland PA14 suspensions carrying each of the three biosensors. The plates were incubated for 24 h at 37°C, and then visualized using the ChemiDoc Imaging System (Bio-Rad, USA) to detect luciferase expression.

### Determination of the minimum inhibitory concentration (MIC)

We assessed the minimum inhibitory concentration of each antibiotic as per the Clinical and Laboratory Standards Institute 2020 protocol [18]. Bacteria grown in LB medium were diluted 100-fold in MH medium and further grown to an OD_600_ of ∼0.1. Then, 10^5^ bacteria from each culture were added to each well of a 96-well plate containing two-fold serial dilutions in medium MH of each of the tested antibiotics, ranging from 16 to 1024 µg/mL in the case of ampicillin; 0.03 to 4 µg/mL in the case of imipenem; 4 to 256 µg/mL for chloramphenicol; 0.5 to 32 µg/mL for tetracycline; 0.007 to 2.5 µg/mL for ciprofloxacin. The plates were then incubated at 37ºC for 24 h. The MIC was obtained by determining the lowest antibiotic concentration that prevented bacterial growth in the plate wells [19, 20].

### Quantitative determination of luciferase

Quantitative assays of luciferase activity were performed using flat bottom white 96-well plates (Greiner, Cat. No. 655098). These plates are optimized for bioluminescence detection. Each well was filled with 10^5^ bacteria (carrying the biosensor) suspended in MH medium containing two-fold serial dilutions of antibiotics, as described for the MIC determination. To quantify luciferase activity the plates were placed in the Synergy H1 multimode microplate reader (BioTek, North Macedonia) at 37°C with stirring, in which both luminescence and OD_600_ were recorded every hour for 18-20 hours. As a positive control, wells filled with bacteria in the absence of antibiotics were added. Luminescence values were normalized against bacterial density in each well (Luminescence/OD_600_).

## RESULTS

### Susceptibility of *P. aeruginosa* to selected antibiotics

To determine the antibiotic susceptibility profile of PA14, a Kirby-Bauer disk diffusion assay with 38 antibiotic disks of different classes (see Materials and Methods) was performed. Figure **1** shows the antibiotics to which PA14 exhibits resistance: ampicillin, cefuroxime, cefoxitin and penicillin G that are associated with cell wall synthesis inhibition; clindamycin that inhibits protein synthesis; trimethoprim, an inhibitor of folic acid biosynthesis; rifampicin (RNA synthesis inhibition) and nitrofurantoin, whose exact mode of action is unknown [21, 22]. Sensitivity to gentamicin was also tested, as the biosensor backbone - pUC18T-mini-Tn7T-*lux*-Gm carries a gentamicin resistance gene.

**FIG 1.**
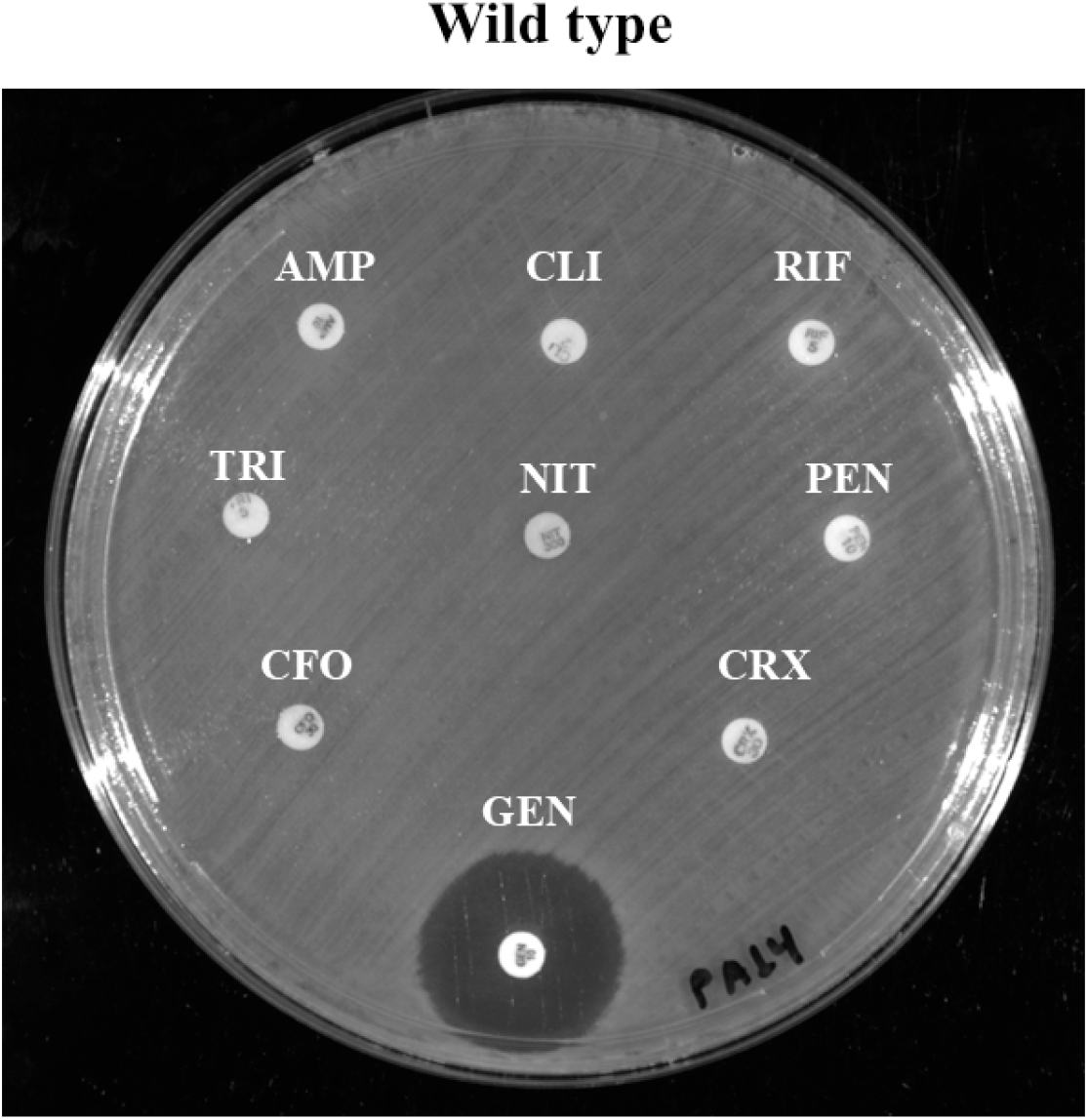
Profile of PA14 resistance to selected antibiotics. AMP (ampicillin); CLI (clindamycin); RIF (rifampicin); TRI (trimethoprim); NIT (nitrofurantoin); PEN (penicillin G); CFO (cefoxitin); CRX (cefuroxime); GEN (gentamicin).

### The ADP-MoA biosensor platform

We developed the ADP-MoA platform as a tool to support the discovery of new antibiotics targeting *P. aeruginosa* while simultaneously identifying their mechanism of action. The platform consists of a set of specific promoters each fused to a promoterless *lux* operon producing thus bioluminescence upon promoter induction. The chosen promoters were: P*recA*, that responds to DNA damage or inhibition of replication [23]; P*ampC*, that as part of the *ampR*-*ampC* system, is activated in response to cell wall damage [11] and P*armZ*, containing a leader peptide-encoding sequence (PA5471.1) upstream of *armZ* whose expression provides a sensor of ribosome function [13]. The amplicons containing the promoter regions were cloned in plasmid pUC18T-mini-Tn7T-*lux*-Gm [16] and transferred to *P. aeruginosa* for integration at the *att*Tn*7* site, located 25 nucleotides downstream of *glmS* [17]. The biosensor strains were evaluated for their responses (luminescence around the growth inhibition halos) to the presence of different antibiotic disks: quinolones (ciprofloxacin, levofloxacin, norfloxacin, ofloxacin, enrofloxacin and nalidixic acid), carbapenems (imipenem, ertapenem and meropenem), penicillins (ampicillin, ampicillin plus sulbactam, ticarcillin, piperacillin plus tazobactam), cephalosporins (cefepime and ceftriaxone), aminoglycosides (amikacin, streptomycin, neomycin, kanamycin, gentamicin and tobramycin) tetracyclines (doxycycline and tetracycline), as well as antibiotics that interact with the ribosome 50S subunit (erythromycin, azithromycin and chloramphenicol) and nitrofuran (nitrofurantoin). As expected, the P*recA*::*lux* biosensor (Figure **2**A) was induced by quinolones and by nitrofurantoin; the P*ampC*::*lux* biosensor (Figure **2**B) responded to a variety of β-lactam antibiotics and the P*armZ*::*lux* biosensor (Figure **2**C) responded to antibiotics that interact with the 30S (aminoglycosides, tetracyclines) or 50S ribosomal subunit (erythromycin, azithromycin and chloramphenicol). Overall, the three biosensors were found to be functional as they responded to all antibiotics of the corresponding class, and thereby could potentially be used for the screening of new antipseudomonal compounds.

**FIG 2.**
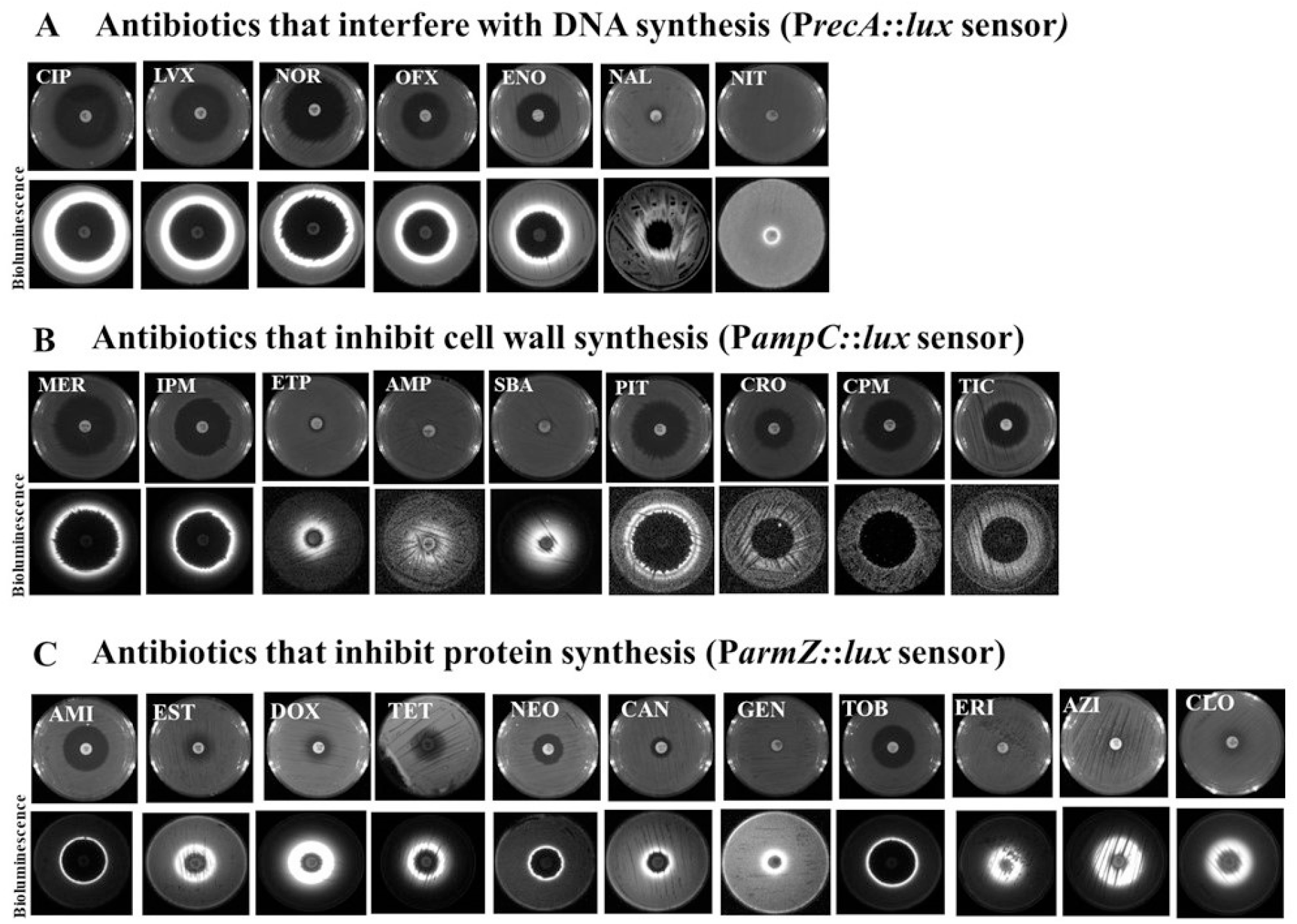
Effect of selected antibiotics on PA14 growth (top) and luciferase activity (bottom) in response to antibiotics that interfere with (A) DNA replication (biosensor P*recA*::*lux*), (B) cell wall synthesis (biosensor P*ampC*::*lux*), and (C) protein synthesis (biosensor P*armZ*::*lux*). The plates were read in the ChemiDoc MP system detector (Bio-Rad, USA) to reveal the luminescence halos. The following antibiotic disks were used: CIP (ciprofloxacin), LVX (levofloxacin), NOR (norfloxacin), OFX (ofloxacin), ENO (enrofloxacin), NAL (nalidixic acid), NIT (nitrofurantoin), MER (meropenem), IPM (imipenem), ETP (ertapenem), AMP (ampicillin), SBA (ampicillin + sulbactam), PIT (piperacillin + tazobactam), CRO (ceftriaxone), CPM (cefepime), TIC (ticarcillin), AMI (amikacin), EST (streptomycin), DOX (doxycycline), TET (tetracycline), NEO (neomycin), CAN (kanamycin), GEM (gentamicin), TOB (tobramycin), ERI (erythromycin), AZI (azithromycin), CLO (chloramphenicol).

### Biosensor specificity

To determine the specificity of the promoter fusions, each biosensor was cross-tested with antibiotics that, according to the literature, should not induce its activity (Figure **3**). As expected, ciprofloxacin induced bioluminescence only in bacteria carrying the P*recA*::*lux* fusion, imipenem only induced the P*ampC*::*lux* promoter and aminoglycosides that inhibit protein synthesis through interaction with the 30S ribosome subunit – amikacin, kanamycin and neomycin, induced only P*armZ*::*lux*. However, chloramphenicol and azithromycin, which interact with the 50S ribosome subunit, induced both P*armZ*::*lux* and P*recA*::*lux*, albeit the latter not as strong as the former. It should be noted that all tested antibiotics inhibited the growth of PA14 irrespective of the constructs they carry – P*recA*::*lux*, P*ampC*::*lux* or P*armZ*::*lux*, except, of course, for the antibiotics shown in Figure **1** and nitrofuran (see below). To further explore the specificity of the biosensors, longer exposition times (> 30 s) were applied to detect faint luminescence signals from weak promoter inductions (see Figure S1 in the supplement). Thirty-three antibiotics were tested: 12 of them affect the cell wall (fosfomycin, bacitracin, ticarcillin, cefepime, ceftriaxone, imipenem, ertapenem, meropenem, piperacillin + tazobactam, ampicillin + sulbactam, clavulanic acid + amoxicillin and ampicillin), 9 inhibit DNA replication (ciprofloxacin, levofloxacin, norfloxacin, ofloxacin, enrofloxacin, nalidixic acid, nitrofurantoin, sulfamethoxazole + trimethoprim and sulfadiazine + trimethoprim), and 11 inhibit protein synthesis (clindamycin, erythromycin, chloramphenicol, azithromycin, doxycycline, tetracycline, kanamycin, streptomycin, amikacin, neomycin and tobramycin). Rifampicin was used here as a negative control (PA14 is resistant to rifampicin, see Figure **1**). Figure S1A shows that with the exception of clindamycin, that did not induce luminescence around the inhibition halo, all antibiotics that interfere with protein synthesis induced P*armZ*::*lux* activity, but not the other biosensors. P*ampC*::*lux*, that responds to cell wall damage, was strongly induced by carbapenems (imipenem, meropenem and ertapenem), ampicillins (ampicillin + sulbactam, ampicillin, ticarcillin, clavulanic acid + amoxicillin, piperacillin + tazobactam) and cephalosporins (cefepime and ceftriaxone), but did not respond to fosfomycin or bacitracin and did not cross-induced other biosensors (Figure S1B). Finally, six antibiotics that affect DNA replication (ciprofloxacin, levofloxacin, norfloxacin, ofloxacin, enrofloxacin and nalidixic acid) induced only the P*recA*::*lux* biosensor. Interestingly, nitrofuran also induced P*recA*::*lux* but did not form a halo of growth inhibition around the disk. Of the antibiotics that affect DNA synthesis, only Sulfamethoxazole + trimethoprim (SUT) were unable to form a luminescent halo (Figure S1C).

**FIG 3.**
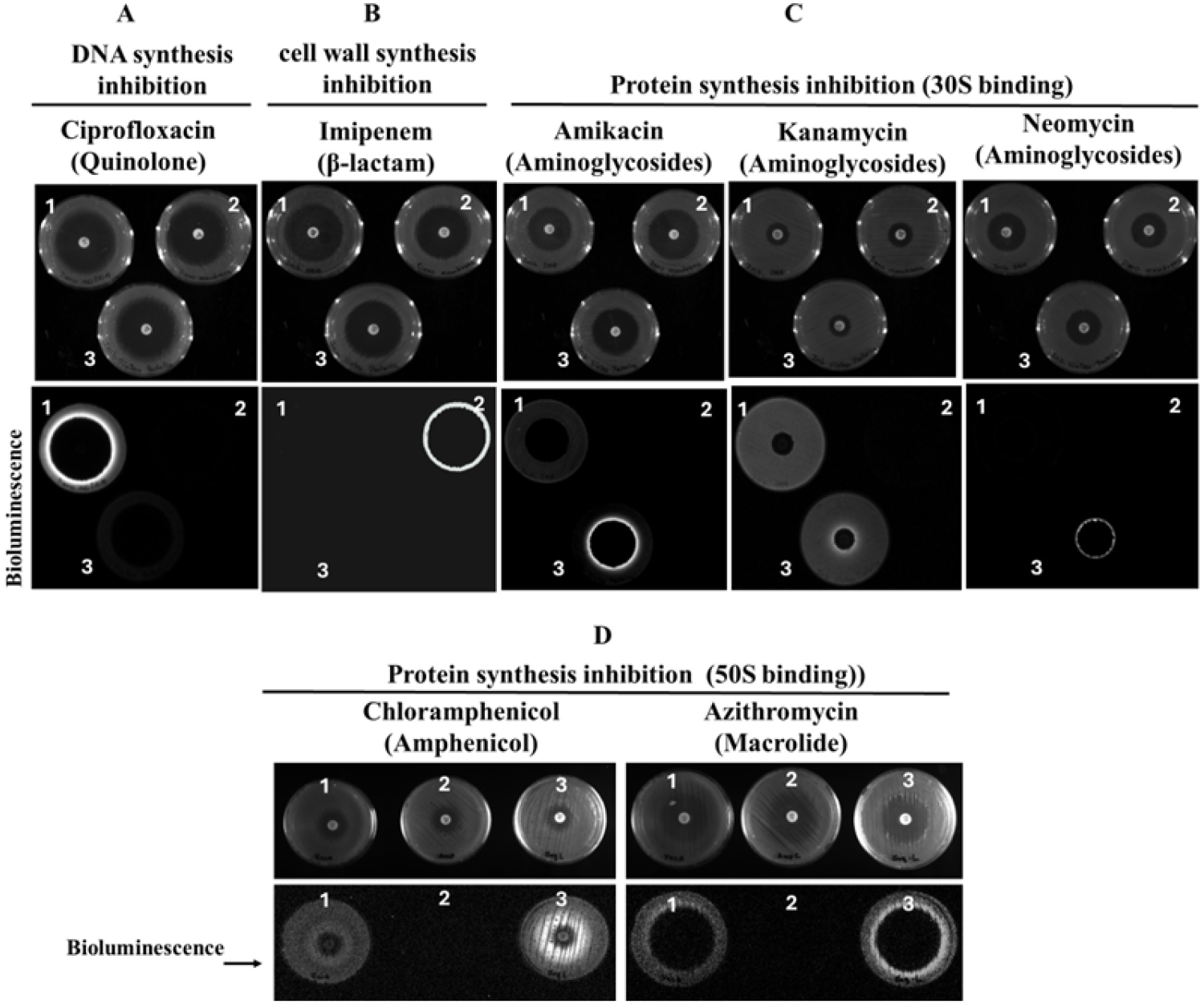
Specificity of the biosensors towards known antibiotics. Bacteria carrying P*recA*::*lux* (1), P*ampC*::*lux* (2) or P*armZ*::*lux* (3) were exposed to disks of each of the following antibiotics: CIP (ciprofloxacin), IPM (imipenem), AMI (amikacin), CAN (kanamycin), NEO (neomycin), CLO (chloramphenicol) or AZI (azithromycin). Images at the top show growth inhibition halos caused by the antibiotics. Images at the bottom show bioluminescence halos elicited by induction of the *lux* fusions. (A) Inhibition of DNA replication by the quinolone ciprofloxacin; (B) inhibition of cell wall synthesis by the β-lactam imipenem; (C) inhibition of protein synthesis by the aminoglycosides amikacin, kanamycin and neomycin; (D) inhibition of protein synthesis by chloramphenicol and azithromycin.

In addition, we confirmed in the present study that the expanded regulatory region of *armZ*, that includes the leader peptide [13] responds to antimicrobials that interfere with protein synthesis, either by interacting with the 30S subunit (aminoglycosides and tetracycline) or with the 50S subunit (erythromycin, azithromycin and chloramphenicol) (Figure **2**, Figure **3** and Figure S1A). Also, this biosensor was weakly induced by some quinolones (Figure S1C).

### Quantification of biosensor response

To obtain a quantitative assessment of the biosensors response, bacteria carrying each of the three constructions (P*recA*::*lux*, P*ampC*::*lux* or P*armZ*::*lux*) were grown in 96-well plates containing MH medium supplemented with subinhibitory concentrations of each of the following antibiotics: ampicillin (64 µg/mL), imipenem (1 µg/mL), tetracycline (4 µg/mL), chloramphenicol (32 µg/mL) or ciprofloxacin (0.03 µg/mL). Bioluminescence was assessed at several point intervals throughout growth (Figure **4**). The time-points varied according to the antibiotic used, some induced earlier responses than others. The P*ampC*::*lux* biosensor was respectively induced by ampicillin and imipenem by more than 4-fold and 20-fold (compared to last time-point before induction) (Figure **4**A and 4B). The other biosensors (P*recA*::*lux* and P*armZ*::*lux*) were not significantly induced by the β-lactam antibiotics. Tetracycline (Figure **4**C) and chloramphenicol (Figure **4**D) specifically induced P*armZ*::*lux* by 86-fold and 60-fold, respectively. P*recA*::*lux* was 12-fold induced by ciprofloxacin (Figure **4**E). Some antibiotics induced the activity of non-related biosensors, but this response was relatively minor. For instance, P*recA*::*lux* activity increased 4-fold in the presence of chloramphenicol (Figure **4**D) and 5-fold in the presence of tetracycline (Figure **4**C) after 17 h of treatment, without further increase after that. P*ampC*::*lux* was induced by 6-fold and 8-fold in the presence of tetracycline and chloramphenicol, respectively, at 17 h treatment onward (Figure **4**C and Figure **4**D). The response to ciprofloxacin was less specific, while P*recA*::*lux* activity was induced by 12-fold, P*ampC*::*lux* and P*armZ*::*lux* were 4-fold induced by this antibiotic (Figure **4**E).

**FIG 4.**
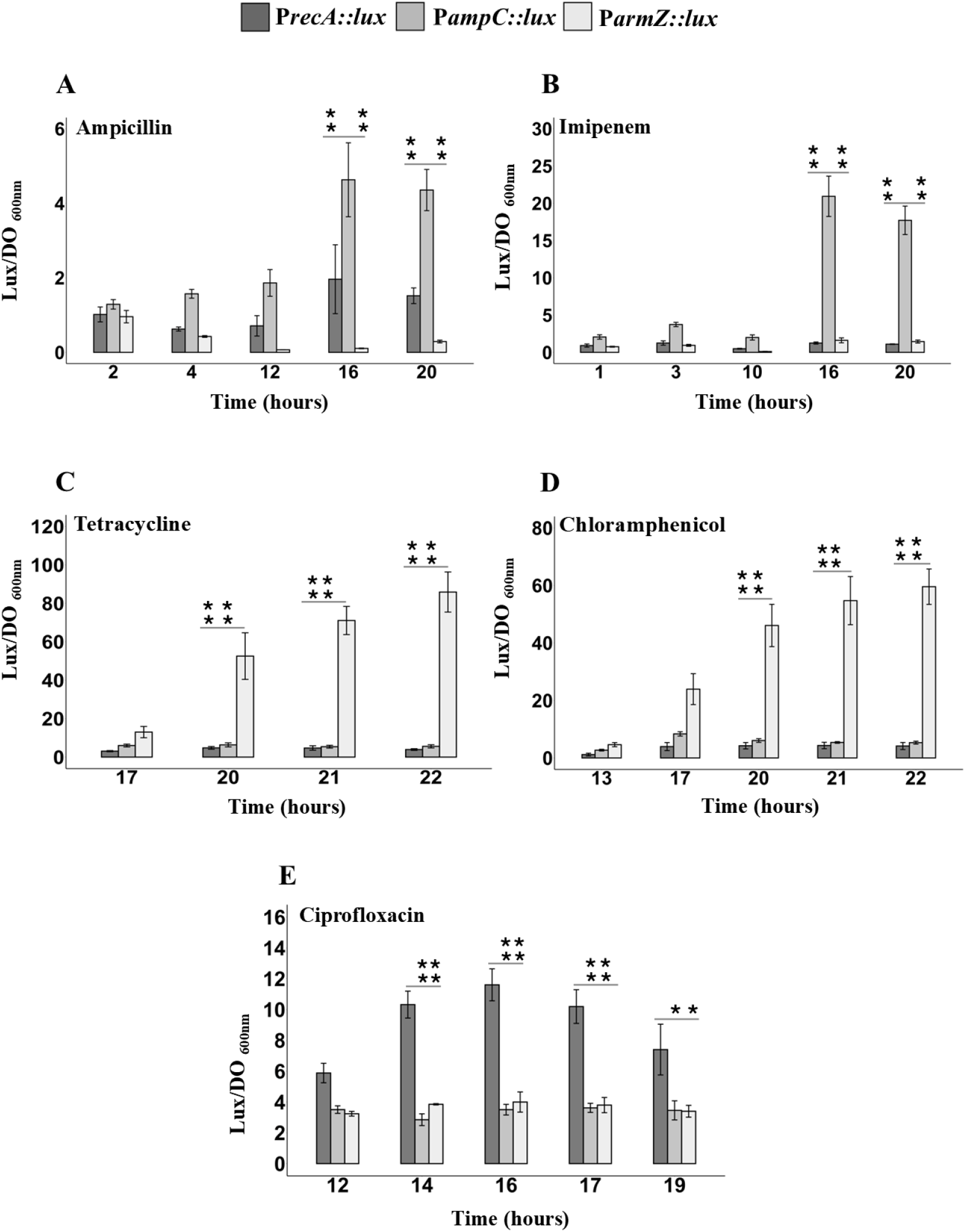
Quantitative analysis of the effect of antibiotics on the biosensors P*recA*::*lux*, P*ampC*::*lux* and P*armZ*::*lux*. Bacteria were exposed to ampicillin (64 µg/mL) (A), imipenem (1 µg/mL) (B), tetracycline (4 µg/mL) (C), chloramphenicol (32 µg/mL) (D) or ciprofloxacin 0,03 µg/mL (E). Luminescence values were normalized by the optical density of the cultures (*lux*/OD_600_) and the results represent the fold-change compared to the control (no antibiotic). ANOVA analysis followed by Tukey: * indicate significant differences (*p* < 0.05); 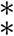 indicates *p* < 0.004 compared with other biosensors. Error bars represent the mean ± standard deviation of three independent biological replicates.

## DISCUSSION

The ADP-MoA platform was designed to aid in the search for potential antibacterial compounds by detecting antibiotic-induced growth inhibition and, simultaneously, identifying the underlying mechanism of action. The platform was hitherto implemented and tested in the PA14 strain of *P. aeruginosa*. We showed that the platform works out in both liquid and solid media and is amenable to high-throughput screening by growing the bacteria in 96 or 384 microplates, providing a low-cost biosensor platform adequate for the screening of molecule libraries of natural or synthetic origin. Table S1 summarizes the effect of all antibiotics tested in this study on the ADP-MoA platform.

The P*ampC*::*lux* biosensor responded to a variety of β-lactams, including carbapenems, penicillins and cephalosporins. All β-lactam antibiotics used in this study activated the expression of *ampC*, including ampicillin, that did not inhibit bacterial growth, but caused a strong bioluminescent response. Carbapenems are resistant to most β-lactamases including *ampC* and extended-spectrum β-lactamases (ESBL) [24]. A key factor in the efficacy of carbapenems is their ability to bind different PBPs, in particular PBP2, PBP4, and PBP5/6 [25, 26]. Cephalosporins, on the other hand, preferentially target PBP1a and PBP3 and penicillin preferentially interacts with PBP3 and PBP4 [26]. We suspect that the strong effect of the carbapenems meropenem, erthapenem and imipinem on *ampC*::*lux* induction observed here is related to their ability to strongly bind multiple different PBPs.

P*armZ* was used with the aim of obtaining a broad-spectrum sensor that responds to molecules interfering with protein synthesis. The *armZ* gene (also known as PA5471) forms an operon with PA5470, which encodes a putative peptide chain release factor [27]. Just upstream to *armZ*, a small ORF (PA5471.1) encodes a 13 amino acid leader peptide, which is freely expressed in the absence of antibacterial compounds. In the presence of antibiotics, ribosome stalls on the PA5471.1 mRNA resulting in alternate mRNA folding enhancing *armZ* expression [13]. In essence, the PA5471.1 peptide acts as a sensor for ribosomal function, inducing the expression of *armZ* in response to antimicrobials that disrupt ribosome activity. In the absence of specific antibiotics, the leader sequence inhibits the transcription of *armZ* through a mechanism of translation attenuation. The induction of *armZ* by antibiotics requires the presence of the entire 367 bp regulatory region located between the PA5472 and *armZ* genes [13]. We showed that P*armZ*::*lux* was strongly induced by all antibiotics that inhibit protein synthesis irrespective of their specific mechanism. For instance, macrolides (azithromycin and erythromycin) and amphenicol (chloramphenicol) that binds to the 50S ribosomal subunit, as well as tetracyclines (tetracycline, doxycycline) and aminoglycosides (amikacin, neomycin, tobramycin, streptomycin and kanamycin) that bind to the 30S ribosomal subunit all induced P*armZ*.

Recent evidence suggests that in addition to their known mechanism of action macrolides, amphenicol and tetracycline, may respectively interact with the V domain of 23S rRNA [28], inhibit the dissociation of the 70S ribosome [29] and bind to the initiation complex comprising the 70S ribosome linked to P-site tRNAfMet and mRNA [30]. These interactions may explain the strong induction of P*armZ* by these antibiotics. On the other hand, aminoglycosides elicited a mild P*armZ*::*lux* response, possibly because aminoglycosides cause mRNA misreading, but not ribosome arrest [31]. This result demonstrates that our sensor is broad-spectrum for ribosome arrest, irrespective of whether the inhibition occurs at the 30S or 50S subunit. Both chloramphenicol and tetracycline induced P*armZ*::*lux*, as expected, but also activated P*ampC*::*lux* and P*recA*::*lux* though at substantially lower levels. Interestingly, it has been reported that chloramphenicol-derived compounds inhibit the early stage of peptidoglycan biosynthesis in *S. aureus* [32]. Also, the tetracycline chelocardin has been shown to exhibit two mechanisms of action: in sub-inhibitory concentrations it inhibits protein synthesis, while at high concentrations it causes both cell wall damage and protein synthesis inhibition [33]. These data might explain why P*ampC*::*lux* is slightly induced in response to these antibiotics.

The P*recA*::*lux* biosensor was induced by all tested quinolones, including nitrofurantoin, whose exact mode of action is not entirely understood [22]. It has been shown that nitrofurantoin inhibits DNA replication by inducing *recA*, which in turn activates the SOS response [34, 35]. Ciprofloxacin induces the SOS response by interfering with the enzymes DNA gyrase or topoisomerase. Interaction with ciprofloxacin generates double-strand breaks (DSBs), resulting in single-strand DNA (ssDNA), which also induces the SOS response [36, 37]. In addition, ciprofloxacin has also been shown to increase the levels of intracellular ROS [38]. Here, we confirmed that ciprofloxacin induces *recA*. However, we also observed that this antibiotic induced other biosensors that respond to cell wall damage and ribosome arrest. Accordingly, in addition to DNA replication inhibition and chromosome fragmentation [39], fluoroquinolones have been shown to cause cytoplasmic condensation by damaging the membrane, leading to cytoplasmic leakage [40]. Another study reported that norfloxacin and ciprofloxacin treatment resulted in some degree of membrane damage [41]. Metabolomics [42] and metabolomic-proteomic [43] studies have identified alterations in transcription, translation, and cell wall synthesis as part of the ciprofloxacin mechanism of action against *Mycobacterium tuberculosis* [42, 44] and *E. coli* [43]. Collectively, these studies suggest that quinolones may have a secondary mechanism of action related to bacteria wall disruption, which might explain the induction of P*ampC* by these antibiotics. In any case, our findings confirm that quinolones operate mainly through *recA* induction.

Finally, the P*recA*::*lux* sensor was slightly induced by chloramphenicol and tetracycline. As mentioned above, some antibiotics in addition of interacting with their specific target, generate ROS that contribute to cell killing [45, 46, 47]. Free radicals also damage DNA [48] and consequently activate the SOS system via *recA* [49].

The use of promoter fusions with fluorescence or luminescence-encoding genes to study the interplay between drugs and gene expression under different conditions is well-established. For instance, Bollenbach et al. [50] analyze a library of ∼110 different *E. coli* promoters fused to the green fluorescent protein (GFP), showing that ribosomal genes are not directly regulated by DNA stress, leading to an imbalance between cellular DNA and protein content. Similarly, Elad et al. [51] evaluated the toxicity and mechanism of action of 420 FDA-approved drugs using 15 different bacterial reporters fused to the *lux* operon, associated with oxidative stress, DNA damage, heat shock, and surplus metal efflux. Valencia et al. [52] used a DNA fusion between the *recA* promoter and *lux* in *P. aeruginosa* PAO1 to show that amikacin prevents ciprofloxacin induction of the SOS regulon. Also, Higuera-Llantén et al. [53] used the concept of biosensors to identify the mechanism of action of new antibiotics.

Our set of biosensors display several improvements: a stable single chromosomal copy of the fusion, which result in a more reliable response by reducing cell-to-cell variability and eliminates the requirement of antibiotics to maintain the plasmid [54]. Another important feature is the fact that in our system the promoters used in the biosensors are derived from the same species and strain as the host – *P. aeruginosa* PA14, which is pathogenic bacterium of clinical interest. In addition, our platform showed a great sensitivity as it allowed the detection of antibacterial compounds that did not cause growth inhibition, as was the case of ampicillin and nitrofurantoin. Another advantage of this platform is that the molecule must penetrate the cell to exert its effect, excluding thus molecules that potentially possess antibacterial activity but cannot go through the membrane barrier. Finally, the ADP-MoA platform can be adapted for the identification of antimicrobials against other pathogens from the ESKAPE group (*Enterococcus faecium, Staphylococcus aureus, Klebsiella pneumoniae, Acinetobacter baumannii, Pseudomonas aeruginosa* and *Enterobacter*), *Mycobacterium tuberculosis*, and others.

### Conclusion

The ADP-MoA platform offers a readily implementable, low-cost approach with significant potential for high-throughput screening of new antimicrobials against *P. aeruginosa* and other bacterial species. The search for new antibiotics is being actively pursued by major projects such as the CARB-X program (https://carb-x.org/) and by the National Institute of Allergy and Infectious Diseases and the National Cancer Institute (USA) [55]. Hopefully, our platform will also be able to contribute to the achievement of this important goal.

## ACKNOWLEDGMENTS

This work was supported by the Conselho Nacional de DesenvolvimentoCientífico e Tecnológico, Brazil (CNPq; grant nº 104461/2019-5) and the São Paulo Research Foundation (FAPESP) PIPE-TC program, process nº 2023/04848-6 and CEPID B3 (process nº 2021/10577-0). Valencia EY was funded by a postdoctoral fellowship from CNPq process nº 104461/2019-5 and fellowship from FAPESP process nº 2024/02524-1. V. Nunes; B. Spira and F. Chambergo are CNPq research fellows.

## DATA AVAILABILITY STATEMENT

Raw data (spreadsheets) are available upon request to the authors.

## AUTHOR BIOGRAPHIES

Author biographies (≤150 words each) and photos (black and white, passport size) can be inserted here. They are required for reviews in CMR and MMBR but optional for minireviews in other journals and Gems in JVI. Do not include them for primary-research articles. See the specific instructions for individual journals for more information.

Use \begin{authorbios}…\end{authorbios} environment to place your author biographies section, commands to do this: output will not come in the pdf for double blind mode.

